# GBMdeconvoluteR accurately infers proportions of neoplastic and immune cell populations from bulk glioblastoma transcriptomics data

**DOI:** 10.1101/2022.11.19.517187

**Authors:** Shoaib Ajaib, Disha Lodha, Steven Pollock, Gemma Hemmings, Martina A. Finetti, Arief Gusnanto, Aruna Chakrabarty, Azzam Ismail, Erica Wilson, Frederick S Varn, Bethany Hunter, Andrew Filby, Asa A. Brockman, David McDonald, Roel GW Verhaak, Rebecca A. Ihrie, Lucy F. Stead

## Abstract

The biological and clinical impact of neoplastic and immune cell type ratios in the glioblastoma (GBM) tumour microenvironment is being realised. Characterising and quantifying cell types within GBMs at scale will facilitate a better understanding of the association between the cellular landscape and tumour phenotypes or clinical correlates. This study aimed to develop a tool that can deconvolute immune and neoplastic cells within the GBM tumour microenvironment from bulk RNA sequencing data. We developed an IDH wild-type (IDHwt) GBM specific single immune cell reference dataset, from four independent studies, consisting of B cells, T cells, NK cells, microglia, tumour associated macrophages, monocytes, mast and DC cells. We used this alongside an existing neoplastic single cell-type dataset consisting of astrocyte-like, oligodendrocyte- and neuronal-progenitor like and mesenchymal GBM cancer cells to create both marker and gene signature matrix-based deconvolution tools. We then applied single-cell resolution imaging mass cytometry (IMC) to ten IDHwt GBM samples, five paired primary and recurrent tumours, in parallel with these tools to determine which performed best. Marker based gene expression deconvolution using GBM tissue specific markers, which we have packaged as GBMdeconvoluteR, gave the most accurate results. The correlation between immune cell quantification by IMC and by GBMdeconvoluteR for primary IDHwt GBM samples was 0.52 (Pearson’s P=7.8×10^−3^) and between neoplastic cell quantification by IMC and by GBMdeconvoluteR was 0.75 (Pearson’s P=1.2×10^−3^). We applied GBMdeconvoluteR to bulk GBM RNAseq data from The Cancer Genome Atlas (TCGA) and were able to recapitulate recent findings from multi-omics single cell studies with regards associations between mesenchymal GBM cancer cells and both lymphoid and myeloid cells. Furthermore, we were able to expand upon this to show that these associations are stronger in patients with worse prognosis. GBMdeconvoluteR is accessible online at https://gbmdeconvoluter.leeds.ac.uk.

**Key points:** GBMdeconvoluteR is a glioblastoma-specific cellular deconvolution tool. When applied to bulk GBM RNAseq data, it accurately quantifies the neoplastic and immune cells in that tumour. It is available online at https://gbmdeconvoluter.leeds.ac.uk

---

Glioblastoma (GBM) brain tumours consist of a multitude of different neoplastic and non-neoplastic cell types[18]. The specific cancer cell subtypes within a GBM are directly influenced by the cellular composition of the microenvironment, which also has a role in shaping the progression of the tumour and its adaption to stressors including treatment[19, 25, 33]. It is of paramount importance to accurately characterise the cellular make-up of GBM tumours. This will enable us to understand the phenotypes associated with changing cell landscapes within individual tumours, and to assess correlation between specific cell populations and the efficacy of new treatments, particularly immunotherapies. Whilst single cell and spatial- profiling approaches currently offer the highest resolution of cellular deconvolution, they are technically challenging, and prohibitively costly for larger sample numbers.

Instead, approaches that propose to quantify cell types from bulk tissue RNA sequencing data have become increasingly popular[2, 4, 16, 20, 24]. These can be split into two main types: those that employ a full cell-type gene expression signature matrix; and those based on marker genes for specific cell types. A widely-adopted implementation of the former approach is CIBERSORTx[20], which was recently used to delineate pan-glioma cell types[33]. However, key studies have shown that the accuracy of any gene expression-based computational deconvolution tool is mostly derived from the signature matrix, or marker genes, underpinning it, which must be derived from the tissue of interest[3, 24, 29]. We, thus, decided to create a tool that can specifically quantify cancer cell types, as delineated by Neftel et al[19], and immune cell types from bulk IDHwt GBM tumour sequencing data. We developed this tool by amalgamating four independent single-cell GBM datasets to derive signature matrices for use with CIBERSORTx and marker genes for use with MCPcounter. The latter was chosen as it has been benchmarked as one of the most accurate marker-gene based tools available, giving consistently high correlation with ground truths across cell types[31]. We then compared results from these GBM-specific programmes to those from orthogonal cell quantification, using single cell-resolution imaging mass cytometry, on the same IDHwt GBM samples. We included both primary and recurrent GBM samples in our tool development and validation, to enable separate quantification of accuracy in longitudinal samples. We found that the MCPcounter based tool performed best at delineating both immune and neoplastic cancer cell populations and have made this publicly available as GBMdeconvoluteR: an online tool accessible via https://gbmdeconvoluter.leeds.ac.uk

## Materials and Methods

All statistical analyses were carried out using the R statistical software package version 4.2.0. The name of each test used, and level of significance achieved, is included within the results where the finding from each hypothesis test is confirmed. Plotting was done using ggplot2 (version 3.3.6).

### Dataset Selection

Four single cell datasets were identified from literature searches (**Table 1**)[5, 23, 28, 35]. The inclusion criteria were single-cell or single-nuclei RNAseq expression data from human IDHwt glioblastoma samples. Data had to be available as raw counts.

**Table 1.** Single-cell *IDH1* wildtype GBM datasets used as a reference set for this project.

| Accession | Samples | Platform | Reference |
| --- | --- | --- | --- |
| GSE141383 | Single cell RNAseq of ~18k cells from 5 primary IDHwt GBM | Automated microwell array capture and full length mRNAseq | [5] |
| GSE163120 | Single cell RNAseq of ~21k cells from primary and ~43kcells from recurrent IDHwt GBMs | 10X Genomics GemCode capture and 3' or 5' mRNAseq | [23] |
| GSE135437 | Single cell RNAseq of 769 cells from 4 IDHwt GBMs | Single cell sorting and 3' mRNAseq | [28] |
| GSE138794 | Single-cell/nuclei RNA-sequencing of ~11k single cells from 4 IDHwt primary GBMs. | 10X Genomics Chromium capture and 3' mRNAseq | [35] |

### Single-cell RNA-seq Data Preprocessing

The Seurat R package (version 4.1.1) was used for all pre-processing, integration, clustering, and annotation tasks[12]. Whilst GSE163120 has a single accession code, it contains data from primary and recurrent sample cells that were sequenced on different platforms so these were processed separately.

#### Copy-number variant analysis to remove neoplastic cells

Single cell datasets were amalgamated. Neoplastic cells were filtered, as has been done previously, by inferring and removing those with large-scale copy number variations such as Chr. 7 amplification and Chr. 10 deletion using the inferCNV R package (version 1.3.3)[1, 21]. The inferCNV object was created using “CreateInfercnvObject()” taking the raw counts (stored in the “RNA” assay of the Seurat object) for each dataset. Annotations were not provided, instead each dataset was grouped according to sample (i.e. patient). The gene ordering file used was derived using the annotations from Ensembl Genes 91 for Human build 38 (GRCh38), taking the gene name, chromosome, and gene span. The “ref_group_names” argument was set to NULL, to average signal across all cells to define the baseline. The “run()” function was then used to perform InferCNV operations to reveal the copy number variation signal. A cut-off value of 1 was used for all the datasets apart from GSE163120, where a value of 0.1 was used as suggested by the documentation for InferCNV.

#### Quality control filtering

Each dataset underwent individual quality control (QC) in which metrics were used to filter out poor quality cells according to dataset-determined thresholds (**Table S1)**: the number of reads, or unique molecular identifiers (nUMI_min); the number of non-zero count genes (nGene); the percentage of mitochondrial genes (mitochondial_ratio_min); the percentage of ribosomal genes; and the cell complexity (gene_complexity_min), which is a composite measure derived as log_10_(nGene)/log_10_(nUMI_min).

#### Dataset normalization

Post-filtering, each dataset was normalised individually using SCTransform, whilst regressing out dataset-specific confounding sources of variation such as ribosomal/mitochondrial ratio using the *vars.to.regress* function argument. Moreover, due to the disparity in the total number of cells in each dataset, a different number of variable features were passed to the *variable.features.n* function argument. The specific normalisation criteria for each dataset are in **Table S2**.

### Dataset Integration

The FindIntegrationAnchors tool was applied to the list of SCTransform normalised datasets to identify cross-dataset pairs of cells that were in a matched biological state. These ‘anchors’ were then used with IntegrateData to merge all the datasets together[12]. The *normalization.method* argument was set as “SCT” for both FindIntegrationAnchors and IntegrateData.

### Clustering and Cell Type Assignment

Dimensionally reduction was performed on the integrated datasets using principal component analysis (PCA) using RunPCA with default settings. This was followed by uniform manifold approximation and projection (UMAP) which was implemented using RunUMAP with custom parameters a=0.6 and b=0.75. Shared nearest-neighbour graphs were constructed based on Euclidean distance using FindNeighbours; taking the default k (k=20), the first 30 principal components and using the *rann* method for finding nearest neighbours. Clusters were identified using FindClusters, with the “smart local moving” (SLM) algorithm used for cluster optimization[34]. The resolution parameter, which sets the ‘granularity’ of the downstream clustering, with increased values leading to a greater number of clusters, was run over a range, in 0.1 increments, between 0.1 – 0.8. The maximum of 0.8 was determined by assessing the best user-defined maximum resolution parameter, based on cluster robustness and stability^21^.

### Cell type annotation

Cell counts per cluster, for each clustering resolution parameter (0.1-0.8) were cross tabulated with immune cell type labels transferred from dataset GSE163120. The 0.7 resolution cross-tabulation (**Table S3**) was used to assign cell-type annotation labels to clusters where the majority of cells had labels for either one distinct cell type or and/or where the cells were labelled were unknown. The T cell, NK cell and TAM labelled clusters could not be assigned and were subclustered to further resolve them. This constituting isolation of these cells and repeat of the above methodology to separate cell types.

### Deriving GBM Immune and Neoplastic Cell Type Profiles

Immune cell marker genes were identified from the integrated, clustered and annotated data using the scran R package (version 1.2.2)[17]. The findMarkers function was used to identify candidate marker genes by testing for those that were differentially expressed (DE) between pairs of clusters using both t-test and Wilcoxon rank sum tests. Both “all” and “any” *pval.type* arguments were used to identify genes which were DE between any two clusters and highly ranked/significantly upregulated genes for a given cluster (all) or significantly upregulated compared with all other clusters (any). The multiMarkerStats function was then used to combine multiple sets of marker statistics. Neoplastic GBM cell marker genes were taken directly from Neftel et al[19] but were filtered to remove non GBM tumour intrinsic genes, to negate the noise that would result from expression of these in the tumour microenvironment[37]. Marker genes for a variety of GBM neoplastic and non-neoplastic cell types have recently been made available as a resource entitled GBMap. We downloaded these directly from the supplementary data of the accompanying preprint for testing within MCPcounter (denoted MCPcounter_GBMap) [. The neoplastic cell markers from GBMap were also filtered to only include GBM tumour intrinsic genes.

### CIBERSORTx reference expression profile

The single cell data used to derive the neoplastic expression profiles used with CIBERSORTx was obtained from the Gene Expression Omnibus (GEO: GSE131928). These data comprised ~23,000 cells which were filtered to include only adult GBM samples. Each cell came with a score corresponding to 6 neoplastic cell states: these were converted to four states and then each cell was assigned to a neoplastic cell state or as a hybrid as described in Neftel et al^2^. The neoplastic single cell data was combined with the labelled immune single cells and then randomly down-sampled such that the total number of cells in the resulting reference matrix was 5075 and of roughly equal class type (**Table 2**)[30].

**Table 2.** Cell types and number of each single cell profile input to CIBERSORTx to develop the GBM specific signature matrix.

| Cell Type | Number of Cells |
| --- | --- |
| AC | 458 |
| B cells | 458 |
| DC | 458 |
| Mast cells | 88 |
| MES | 458 |
| Microglia | 458 |
| Monocytes | 458 |
| NK cells | 458 |
| NPC | 458 |
| OPC | 407 |
| T cells | 458 |
| TAM | 458 |
| **TOTAL** | **5075** |

### Validation Samples

Ten human GBM samples were used for validation via bulk RNA sequencing and imaging mass cytometry. These were *de novo* primary IDHwt GBM that had been stored in formalin-fixed, paraffin-embedded blocks, and the matched locally recurrent sample following initial debulking surgery and treatment with radiation and Temozolomide chemotherapy.

### Ethics Statement

Samples were from patients at the Walton Centre, UK, that provided informed consent in writing for the use of their tissue in research. The inclusion of these samples in this project was following approval by the UK National Health Service’s Research Ethics Service Committee South Central - Oxford A (Research Ethics Code: 13/SC/0509).

### Bulk RNA sequencing

RNA was extracted from neuropathologist annotated regions containing >60% cancer cells using Qiagen kits (Qiagen, Sussex, UK). Paired end, 100bp strand-specific whole transcriptome libraries were prepared using the NEBNext Ultra Directional RNA Library Prep Kit for Illumina (New England BioLabs, Herfordshire, UK), following rRNA depletion with NEBNext rRNA Depletion Kit or Ribo-Zero Gold. Libraries were sequenced on an Illumina NextSeq2000. RNAseq data was processed as previously described[9].

### Imaging Mass Cytometry (IMC)

#### Antibody Selection

A panel of 33 antibodies for markers of neoplastic and immune cell subtypes in GBM was selected based on literature searches and manufacturer websites as collated in **Table 3** and **Table S4**. Neoplastic GBM cell markers were selected based on an overlap of GBM cancer cell delineators from three independent, single cell studies, including the Neftel et al. study that underpins the gene expression approach herein[7, 19, 35]. Antibody selection criteria was, in order of priority: available in pre-conjugated format for IMC and previously used in IMC of GBM or normal brain; previously used in IMC of GBM or normal brain via bespoke conjugation; available in carrier free format and had been validated for use in IHC or ICC in brain or GBM; available in carrier free format.

**Table 3.** Antibodies used in IMC.

| Marker/Target | Cell Type | Functional State/specifics | Antibody clone(s) |
| --- | --- | --- | --- |
| ANXA1 | GBM cancer cells | Hypoxia driven mesenchymal | EPR19342/abcam |
| ANXA2 | GBM cancer cells | Hypoxia driven mesenchymal | MAB3928/RnD |
| BCAN | GBM cancer cells | Neural progenitor like | S294A-6/Thermo |
| CD3 | Immune cells | T-cells | Fluidigm/3170019D |
| CD31 | Normal brain cells | Vasculature | Fluidigm/EPR3094 |
| CD45 | Immune cells | Pan-immune marker | Fluidigm/3152016D |
| CD8 | Immune cells | T-cells | SK1/Biolegend |
| CHI3L1 | GBM cancer cells | Mesenchymal | EPR19078-157/abcam |
| DNA | All cells | Cell nucleus | Fluidigm |
| DLL3 | GBM cancer cells | Neural progenitor like | EPR22592-18/abcam |
| EZH2 | All cells | Chromatin remodeller | EPR9307(2)/abcam |
| GFAP | Normal brain cells | Astrocyte | ab218309 /abcam |
| HIF1A | All cells | Hypoxia | 16H4L13/Thermo |
| HOPX | GBM cancer cells | Astrocyte like | ab230544 |
| IBA1 | Immune cells | Pan-macrophage | EPR16588 /abcam |
| JARID2 | All cells | Chromatin remodeller | Developed in house |
| JARID2 | All cells | Chromatin remodeller | EPR6357/abcam |
| Ki67 | All cells | Proliferating cells | B56/Fluidigm |
| MOG | Normal brain cells | Oligodendrocytes | MA5-24644/Thermo |
| NCAM (CD56) | Normal brain cells | Immature Neuron | HCD56/Biolegend |
| NeuN | Normal brain cells | Mature Neuron | 1B7/Biolegend |
| NKp46 | Immune cells | NK cells | MAB1850/RnD systems |
| OLIG1 | GBM cancer cells | Oligodendrocyte progenitor like | MAB2417/R&D |
| P2Y12R | Immune cells | Microglia | EPR23511-72/abcam |
| SCD5 | GBM cancer cells | Oligodendrocyte progenitor like | PA5-59963/Thermo |
| SLC1A3 | GBM cancer cells | Astrocyte like | EPR12686/abcam |
| SMA | Normal brain cells | Vasculature | 1A4/R&D |
| SNAI1 | GBM cancer cells | Epithelial to Mesenchymal Transition | AF3639/R&D |
| SOD2 | GBM cancer cells | Mesenchymal | EPR2560Y/abcam |
| SOX2 | GBM cancer cells | GBM stem-like cell | O30-678/Fluidigm |
| TGFbeta | GBM cancer cells | GBM stem-like cell | TW4-6H10/Fluidigm |
| TMEM119 | Immune cells | Microglia | HPA051870/sigma |
| TNC | GBM cancer cells | GBM stem-like cell | MAB2138/R&D |

A set of panel-wide control tissues was determined: spleen, brain, tonsil, prostate, bone marrow, skin and uterus. Control tissue samples from at least two individuals were amalgamated into a multi-tissue formalin fixed, paraffin embedded block. Multi-tissue block sections were used in IHC validation and testing of three antibody concentrations at, above and below those recommended by the manufacturer. Chosen antibody concentrations and control tissue(s) relevant to each antibody are in **Table S4**. Antibody conjugation and staining and IMC took place at the Flow Cytometry Core Facility at Newcastle University. Conjugation was performed using MaxPar metal labelling kits using X8 polymer according to standard manufacturers protocols (with the exception of Gd157 which was obtained by Trace Sciences International and was diluted to 0.1M prior to use with MaxPar reagents). Conjugations were validated by capture on Thermo AbC beads prior to acquisition on a helios mass cytometer.

#### Sample Preparation and Mass Cytometry

5μm sections, taken consecutively from the same blocks that underwent bulk RNA sequencing (see above), were stained with a cocktail of all 33 conjugated antibodies after dewaxing (Xylene) and HIER antigen retrieval in Tris-EDTA (pH9) with 0.5% Tween 20. Sections were incubated for 30 minutes in 0.3 uM irridum to counterstain the nuclei prior to air drying. A minimum of three 2mm^2^ regions of interest (ROI) were annotated per sample within the area corresponding to that from which RNA was extracted from the adjacent sections. Images were generated on the Hyperion Tissue Imaging cytometer by ablation of the ROI at a 200Hz frequency with a 1-micron diameter laser. Raw MCD files were created and exported as ome-tiff from MCD Viewer software (Fluidigm).

#### Image Pre-processing

Following export, the raw data were converted from to ome-tiff format and segmented into single cells using the *steinbock* pipeline comprised of the following steps^[38]^. Pixel classification was done using Ilastik (version 1.3.3): Tiff stacks were generated for each of the proteins in the panel and pixels classified into two channels as either nuclear, or background. These were used to train a random forest classifier, which returned probability masks for each image. The generated probability maps were processed to create single-cell masks using the image analysis software CellProfiler (version 4.1.3). First, probabilities were histogram-equalized (256 bins and kernel size of 17), and then a Gaussian filter was applied to enhance contrast and smooth the probabilities. Subsequently, an Otsu two-class thresholding approach was used to segment nuclear masks. Cell masks were derived from an expansion of nuclear masks using a maximum expansion of 3 pixels. The CellProfiler single cell masks were ultimately overlaid onto the single-cell segmentation masks and single-channel tiff images of all measured channels to extract single-cell marker expression means. The single-cell data was read into R using read_steinbock from the imcRtools R package (version 1.2.3) and the expression counts were transformed using an inverse hyperbolic sine function (cofactor = 5). The expression counts were corrected for channel spillover using a non-negative least squares method as previously described[6]. Briefly, each metal-conjugated antibody was spotted on an agarose-coated slide, and this was ablated to generate a background signal which could be used for compensation using the R Bioconductor package CATALYST (version1.20.1).

#### Image Analysis

All downstream data visualisations, including Image and cell segmentation quality control were completed using the cytomapper (version 1.8.0) and dittoseq (version 1.8.1) R packages[10]. Batch effect correction of segmented cells was completed using harmony (version 0.1.0)[14]. Cells were clustered based on their similarity in marker expression using the PhenoGraph clustering algorithm (k =45) implemented in Rphenograph (version 0.99.1)[15]. Cluster IDs were mapped on top of UMAP embeddings (n_neighbors = 40) derived using the uwot R package (version 0.1.11). Cell type classification was completed using marker enrichment modelling, implemented in the MEM R packages (version 2.0.0), selecting for markers with enrichment scores equal to or greater than 3 (display.thresh = 3)[8] for the first clustering, which defined immune cells. Further subclustering was required to annotate neoplastic cells with display.thresh relaxed to 2 (**Table S5**).

### Creating and Comparing the Cell Deconvolution Approaches

MCPcounter was run via the R Package (version 1.2.0) in two modes: default mode(MCP_default_) used the universal set of 110 immune cell-type marker genes that come provided as standard, meaning no neoplastic cell populations were included; GBM mode (MCP_GBM_) used the GBM-specific neoplastic and immune cell marker genes derived as outlined above.

The ‘Create Signature Matrix’ module of CIBERSORTx was run with default parameters and quantile normalization disabled, to create a signature matrix using the single-cell-derived immune and neoplastic expression profiles detailed above. This signature matrix was then used to infer cell fractions of bulk RNA-Seq sample mixtures using the CIBERSORTx High-Resolution docker container (https://hub.docker.com/r/cibersortx/hires). For all runs, the bulk RNAseq dataset was input as the ‘mixture’ file and the respective signature matrix was input as the ‘sigmatrix’ file. For all runs, the Batch correction was done in ‘S-mode’ by setting the ‘rmbatchSmode’ parameter to TRUE and the input signature matrix’s respective CIBERSORTx-created “source gene expression profile” was input. Finally, absolute mode was set to FALSE for all runs. Cell population quantities inferred from the GBM sample RNAseq for all expression-based deconvolution approaches were compared with those from the IMC using the Pearson Correlation Coefficient.

### Application to TCGA data

TGCA data was obtained from the Genomics Data Commons Data Portal (https://portal.gdc.cancer.gov/). The data were filtered on the “data_category” and “data_type” fields to only include “transcriptome profiling” and “Gene Expression Quantification” data, respectively. Further, only primary, IDH wild-type GBM cases treated with standard/non-standard temozolomide chemoradiation were selected. The expression values for the 93 samples that were obtained were unlogged, TPM normalised counts which were combined into an expression matrix that was then input to GBMdeconvoluteR run using our GBM specific marker genes. Outputted scores were used in correlation analysis using the cor() and cor.test() functions from base R stats package. The quartiles of overall survival (OS) were calculated and used to extract patients with a worse (OS less than the lower quartile of 8.55 months) or better (OS greater than the lower quartile of 20.55 months) prognosis. Plots were generated using the ggplot2 R package.

### Developing GBMdeconvoluteR

GBMdeconvoluteR was developed as an interactive web application using the Shiny R package (version 1.7.1) and packaged as a portable container image using the rocker/shiny:latest base Docker image. The custom image was stored in the Azure Container Registry and deployed using the Azure App Service. All code can be found at https://github.com/GliomaGenomics/GBMDeconvoluteR.

## Results

### Identifying GBM Specific Cell Type Profiles

Four independent single cell GBM datasets (**Table 1)** were used to derive marker genes, or signature gene expression matrices, for GBM tumour-infiltrating immune cells: B cells, T cells, natural killer (NK) cells, microglia, tumour associated macrophages (TAM), monocytes, mast and dendritic cells (DC). **Figure 1A** outlines the process. Datasets underwent pre-processing independently to filter out poor quality cells and copy number analysis to remove neoplastic cells, before being amalgamated. There were significant batch effects owing to different sequencing platforms and originating centres but these were effectively removed using regularized negative binomial regression[11] (**Figures 1B and S1A**). One dataset (GSE163120) included the immune cell annotations determined by the original study. This information was used to guide clustering, with optimisation focused first on maximising cluster stability and then on the best separation of pre-annotated cell types[22]. Owing to the difficulty in separating immune types that are known to have similar and overlapping gene expression profiles (namely tumour associated macrophages [TAM] and microglia; and natural killer [NK] and T-cells) cells assigned to any of these groupings were isolated and further sub-clustered, resulting in definitive cluster annotations (**Figure 1B and S1B**).

**Figure 1.**
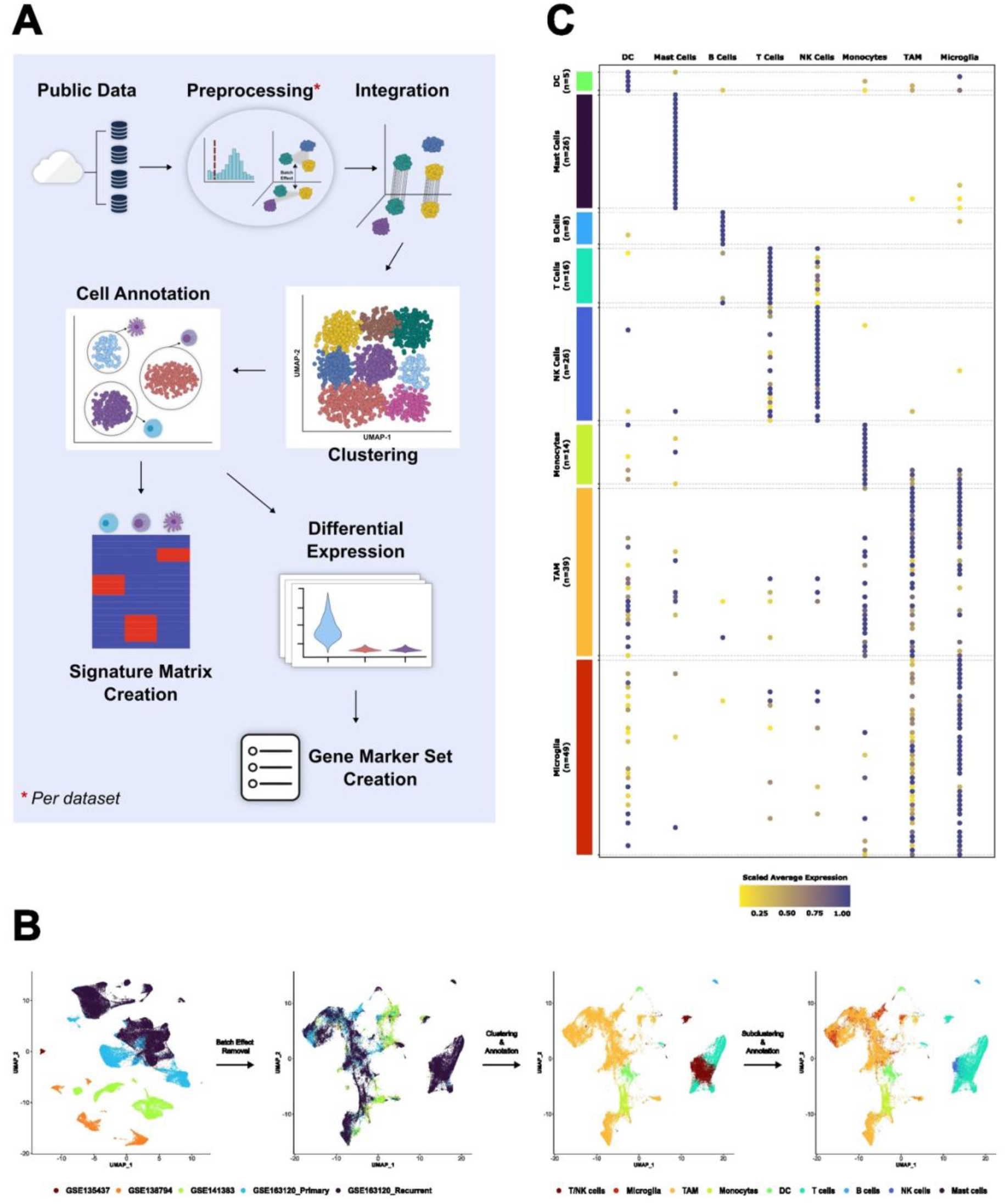
**A**. The process adopted to amalgamate several independent single cell GBM datasets and create a GBM-specific immune cell reference signature gene expression matrix (for input to CIBERSORTx) or marker gene set (for input to MCPcounter). **B**. The inherent batch effects in the amalgamated data are evident in dimensionality reduction plots where clusters initially separated by originating datasets (far left), but were removed by normalisation (middle left and Figure S1A). Initial clustering and cell type assignment of the normalised data was unable to resolve TAM and microglia, and T- and NK-cells (middle right) but further sub-clustering enabled these cell types to be further delineated (far right and Figure S1B). **C**. A dot plot showing the expression of chosen GBM-specific immune cell type markers (y-axis) in each cell type in the amalgamated single cell data (x-axis).

GBM-specific marker genes for each immune cell type were then derived by using differential expression analysis to highlight the top 25 genes, per annotated cluster, that were uniquely or predominantly expressed in that cluster, and visually checking these to identify specific cell type markers corresponding to each immune cell type (**Figure 1C** and **Table S6)**. Marker genes for GBM cancer cell subtypes were adopted from Neftel et al[19]. In that study, four neoplastic GBM cell types were delineated from single cell data. We extracted the marker genes that Neftel et al. showed to delineate the four subtypes, but then removed those that are also expressed in the GBM tumour microenvironment, and would therefore confound the results of application to bulk tissue profiles[37] (**Table S7)**.

Single cell expression profiles for annotated GBM-associated immune cells, from our combined datasets, or for annotated GBM cancer cell subtypes, from Neftel et al., were amalgamated into a full gene expression matrix. This was then subsampled to produce a total of 5075 single cell gene expression profiles with roughly equal representation of each cell type (**Table 2**).

### Developing and Validating the Deconvolution Approach

Two gene-expression based computational deconvolution approaches were investigated owing to previous benchmarking studies finding them to be the best full gene expression signature matrix-based approach (CIBERSORTx) and marker-gene based approach (MCPcounter) available[31]. The approaches are distinct and give results with different interpretations. Gene expression signature matrix methods such as CIBERSORTx attempt to quantify cell types in a single sample, enabling comparison of proportions of all cell types within and between samples. Marker gene based methods like MCPcounter instead score a single cell type for comparison of prevalence across samples; the score from cell type A cannot be compared with cell type B so within sample comparisons of different cell types is not possible. To ascertain the accuracy of these programmes and determine which performed best, we identified five primary and matched recurrent GBM samples on which to perform both gene expression-based and imaging mass cytometry (IMC)-based cell type deconvolution (**Figure 2A and Figure S2**). The latter is an approach that characterises cells, according to protein expression, at single cell resolution in tissues using up to 40 antibodies (**Figure 2B**). We assembled and validated a panel of antibodies known to distinguish tumour-infiltrating macrophages, microglia, monocytes, NK and T-cells (**Tables 3 and S4**).

**Figure 2.**
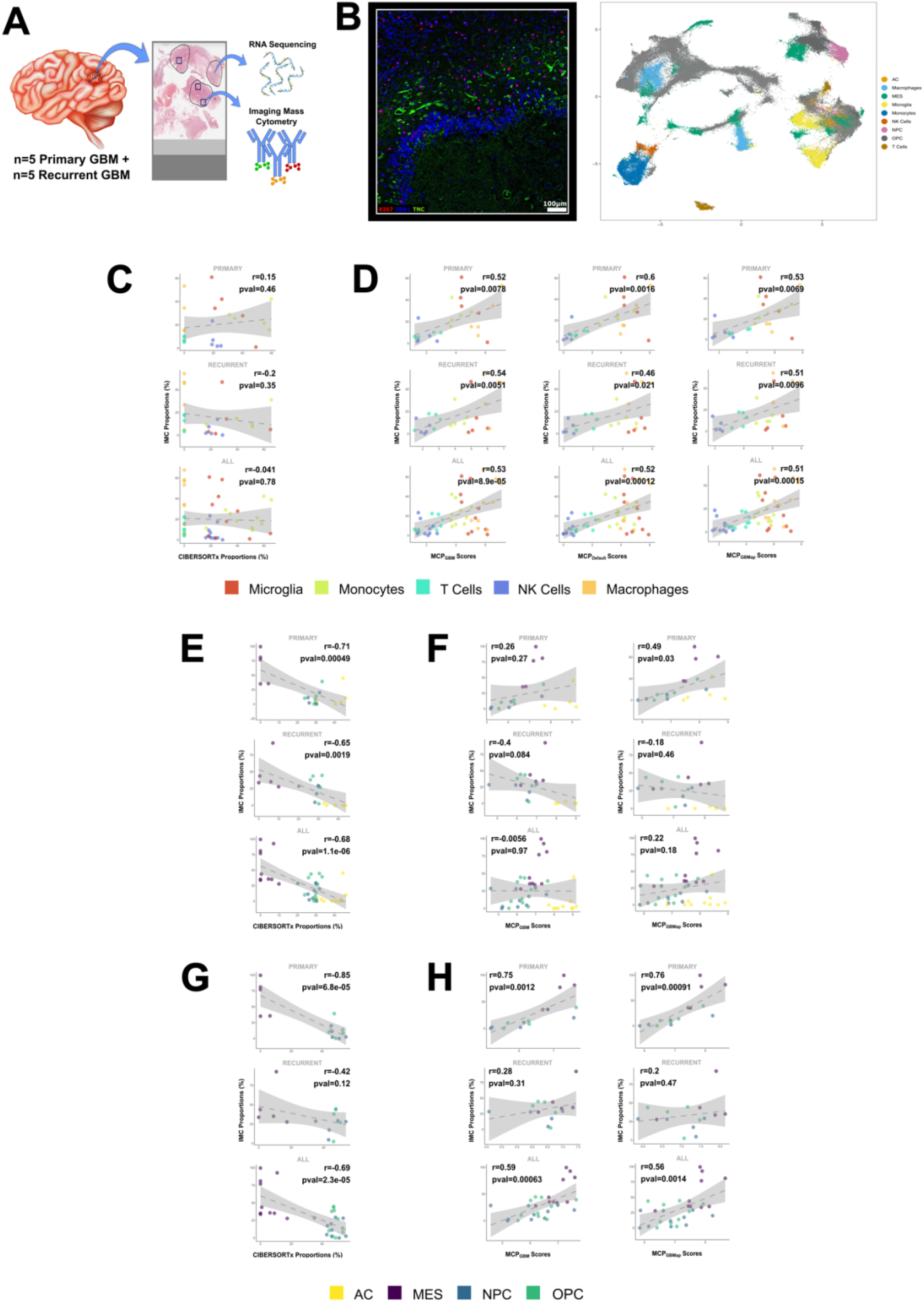
**A**. A schematic showing how patient samples were used for validation. Regions of formalin fixed tissue sections were annotated as high tumour cell content by a neuropathologist (marked in black) and were macro-dissected for RNA sequencing. At least three overlapping regions (blue squares) per sample were subjected to imaging mass cytometry (IMC) on a consecutive section. **B**. Left: A representative image from the IMC for GBM sample 64 with three of the chosen protein markers annotated.Right: The UMAP projection of cell types assigned according to the expression of cell type protein markers quantified by IMC. **C-H**. Scatterplots of gold standard cell proportions quantified by IMC versus CIBERSORTx using a GBM-specific reference gene signature matrix for immune cells **(C)**, and neoplastic cells with **(E)** and without **(G)** astrocyte-like GBM cells included. Scatterplots of cell proportions quantified by IMC versus the cell type relative prevalence score from MCPcounter using different maker gene sets for immune cells (**D)**, and neoplastic cells with **(F)** and without **(H)** astrocyte-like GBM cells included. Marker genes for MCPcounter were either default (MCP_default_), GBM-specific according to our research (MCP_GBM_) or GBM-specific according to GBMap (MCP_GBMap_) Neoplastic cells are astrocyte-like (AC), oligodendrocyte progenitor-like (OPC), neuronal progenitor-like (OPC) or mesenchymal (MES). Pearson’s correlation coefficients (r) and p-values are indicated on each plot.

#### Immune Cell Proportions

We first inspected the concordance between immune cell proportions predicted by CIBERSORTx and quantified by IMC, as the ground truth, for primary and recurrent GBM tumours separately and then all tumours together (**Figure 2C**). The performance in primary samples (Pearson’s r=0.15) was better than for recurrent samples (Pearson’s r=−0.2) but no results were significantly correlated (Pearson’s p<0.05) and even where positive, correlation coefficients remained low.

#### Immune Cell Prevalence

MCPcounter can be used in default mode in which in-built canonical immune cells markers are employed. When running the programme in this mode it can only be used for immune cell estimation and we refer to it as MCP_default_. In contrast, the mode using the GBM-tissue specific immune and neoplastic cell markers listed in **Tables S6 and S7** is denoted MCP_GBM._ In addition, at the time of preparing this manuscript a larger GBM-specific single cell resource, GBMap, was made available that amalgamated 26 single cell brain and GBM datasets [27]. We, thus, also ran MCPcounter using the GBMap marker genes, denoting this as MCP_GBMap_. We inspected the concordance between the relative cell type prevalence scores that resulted from each version of MCPcounter and the quantification by IMC (**Figure 2D**). All MCPcounter based approaches were more accurate than CIBERSORTx, with MCP_GBM_ performing best over all samples (Pearson’s r and p-values are: 0.53 and 8.9×10^−5^ between MCP_GBM_ and IMC; 0.52 and 1.2×10^−4^ between MCP_default_ and IMC; and 0.51 and 1.5×10^−4^ between MCP_GBMap_ and IMC).

#### Neoplastic Cell Quantification

The four GBM cell types described by Neftel et al. are delineated by gene expression [19]. Recent studies have shown that such transcriptional cell-type markers often do not translate to protein level markers for use in approaches such as IMC[13, 32]. We set out to test this for the GBM neoplastic cell types, specifically. To that end, in our IMC panel we included antibodies against markers of the four neoplastic GBM cell types from Neftel et al., prioritising those that overlapped with markers of GBM cancer cell subsets identified in two independent studies: Wang et al.[35] and Couturier et al.[7] (**Tables 3 and S4; Figure S3**). These studies also identified GBM cancer cell subsets that were labelled differently but showed good agreement with the Neftel et al. study.

Results showed poor concordance over all samples for both gene-signature based (**Figure 2E**; Pearson’s r and p-values are: −0.68 and 1.1×10^−6^ between CIBERSORTx and IMC;) and marker gene based approaches (**Figure 2F;** Pearson’s r and p-values are: −0.0056 and 0.97 between MCP_GBM_ and IMC; 0.22 and 0.18 between MCP_GBMap_ and IMC). However, a closer inspection indicated that a single GBM cancer cell type, the astrocyte (AC)-like cells, were impacting the overall correlation between IMC and the gene expression-based approaches. Removing AC-like cells from the analysis (**Figures 2G-H)** showed strong concordance between RNAseq and IMC for primary GBM samples for both marker based methods (r≧0.75, albeit with weaker concordance for recurrent samples. This suggests that AC-like cells were not being correctly quantified using transcriptionally-delineated markers for IMC, but the remaining three cancer cell types were. Based on this, we proceeded to evaluate the ability of each method to perform relative neoplastic GBM cell type quantification with AC cells removed. We found MCP_GBM_ to be the most accurate over all samples (**Figures 2G-H;** Pearson’s r and p-values are: 0.59 and 6.3×10^−4^ between MCP_GBM_ and IMC; 0.56 and 1.4×10^− 3^ between MCP_GBMap_ and IMC).

### Application to TCGA data

Our results show that MCP_GBM_ is able to accurately quantify immune and neoplastic cells in GBM tissue bulk sequencing data. To show how this can be useful in gaining biological and clinical insights from large-scale studies, we applied MCP_GBM_ to bulk RNAseq data from 93 GBM samples from The Cancer Genomic Atlas (TCGA). This gave a score per cell type per sample, allowing us to quantify the correlation of cell type prevalence across patients (**Figure 3A**). Recent spatial, multi-omics studies have suggested that different neoplastic GBM cell types associate with, and are programmed by, different environmental niches of GBM tumours[25]. A key finding was that mesenchymal (MES) cancer cells associate with both myeloid and lymphoid compartments, whereas the remaining neoplastic cell types (AC-, NPC- and OPC- like cells) are significantly depleted in immune-rich regions. Our results recapitulate these findings: we oberrved significant, high, positive correlations between MES and all immune cells quantified, and significant negative correlations between the remaining neoplastic cell types. This phenomenon was more pronounced for non-MES neoplastic cells associated with neuronal development (NPC- and OPC-like cells) than for AC-like cells, also in keeping with the previous findings[25]. Based on the high numbers of samples in TCGA we were able to further separate patients using overall survival (OS) quartiles to extract worse prognosis (OS less than the lower-quartile of 8.55 months) and better prognosis (OS greater than the upper-quartile of 20.55 months) cohorts and compare score distributions (**Figure 3B**) and correlations (**Figure 3C**) in these patient subsets. The prevalence scores of cell types is not significantly different between worse or better prognosis patients (**Figure 3B**) but the correlations between cell-types are markedly different (**Figure 3C**). Patients with worse prognosis have higher and more significant correlations (both negative and positive) between neoplastic and immune cell types. The tumour microenvironment has been shown to shape the neoplastic cell landscape over time in GBM, with more aggressive tumours being linked to greater polarity and classification of neoplastic subtypes[25, 33, 36]. Our results suggest that, in worse prognosis tumours, neoplastic and immune cells are more tightly associated, potentially through more direct inter-cellular communications, which could be promising therapeutic targets. These preliminary results exemplify how our tool can be used to develop new insights and hypotheses, by being applicable to large scale datasets.

**Figure 3.**
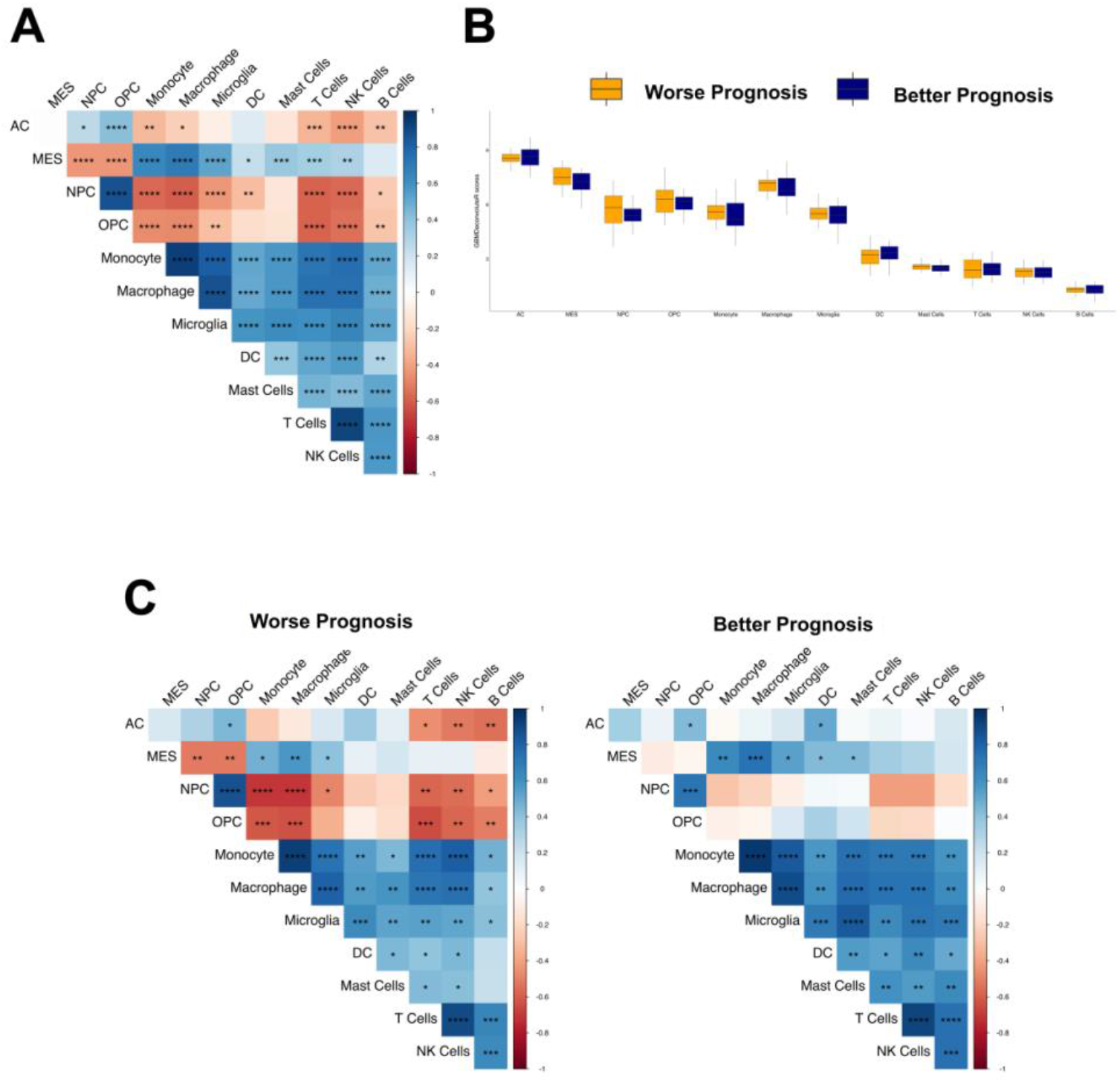
MCP_GBM_ was used to score cell types in bulk GBM RNAseq data from The Cancer Genome Atlas (TCGA). **A**. Heatmap of the correlations between cell type scores across all samples. **B**. Boxplots showing distribution of cell type scores for patients with worse or better prognosis (determined by the lower and upper quartile of overall survival, respectively). **C**. Heatmap of the correlations between cell type scores across samples from patients with worse (left) or better (right) prognosis. Significance is denoted by asterisks: *: p<0.05; **: p<0.01; ***: p<0.001; ****: p<0.0001.

### Incorporating additional GBM cell types and Making Our Approach Available Via GBMdeconvoluteR

To make MCP_GBM_ available to the neuro-oncology community, we have packaged it into an online application called GBMdeconvoluteR. We also give the user the option to use the marker genes from GBMap[27] because, although these did not quantify cell types as accurately as MCP_GBM_, the GBMap reference set extends the range of GBM non-neoplastic cell types that can be quantified from bulk expression data. GBMdeconvoluteR is, thus, a web-based application that enables users to upload bulk GBM expression profiles and output the relative proportion of immune and neoplastic GBM cells, or using GBMap markers genes as input, to also include normal brain and blood-vessel cells, across multiple samples.

## Discussion

We have developed the first publicly available GBM specific deconvolution tool that can infer both neoplastic and non-neoplastic cell population prevalence from bulk GBM tumour RNA sequencing data. This tool was developed by amalgamating four independent, human, single cell sequencing datasets to create tissue specific cell type gene expression reference profiles. The single cell data was from *de novo* IDHwt GBM either at initial diagnosis (primary) or upon recurrence. Recurrent GBMs have been shown to have altered transcriptional profiles which may impact on the accuracy of the deconvolution results[26, 33], so we included these samples in the tool development and validation. We found that our approach is suitable for deconvoluting recurrent GBM tumours but, in keeping with the aforementioned studies, neoplastic cell deconvolution is not as accurate at the longitudinal time point. Our study confirms, as shown elsewhere, that tissue specific reference datasets are necessary to achieve maximal accuracy in expression-based deconvolution[3, 24, 29].

We used imaging mass cytometry (IMC) to ascertain the ground truth of cell type characterisation and quantification. We then compared this with the results from the gene-expression-based approaches to determine which should underpin our tool, and to establish its accuracy. However, it must be noted that the regions that underwent IMC, whilst encompassed within, were substantially smaller than regions that underwent RNAseq (**Figures 2A and S2**), and that the GBM microenvironment is notoriously heterogeneous[25]. That, and the fact that IMC was performed on different, albeit, adjacent tissue sections, means that a deviation from perfect correlation is not just a result of gene expression deconvolution tool performance, but also in bona fide differences in cell proportions.

Our study is the first to evaluate whether the marker genes of the four GBM neoplastic cell types, determined by Neftel at al. from gene expression data, are preferentially expressed at the protein level. We found that for MES, NPC and OPC GBM cells there was a clear association between the protein levels of the selected markers and the gene expression-based quantification, but this was not the case for the markers chosen to proteomically identify AC-like cancer cells (SLC1A3 and HOPX). This further highlights the need to validate cell type markers identified either at the protein or gene expression level, prior to use via the other modality.

GBMdeconvoluteR is a publicly available webserver, enabling researchers to accurately determine the cell types and prevalence in GBM samples from bulk RNAseq data. The marker-gene MCPcounter based method was the most accurate. It should be noted that marker-based deconvolution results in relative, rather than absolute, cell type quantification meaning comparison is possible within cell types across samples, rather than within samples across cell types. We applied GBMdeconvoluteR to data from TCGA and were able to confirm recent findings from single cell resolution multi-omics studies, regarding the specific enrichment of MES neoplastic cells in immune compartments, and depletion of other GBM cancer cell types. However, because our approach is easily applicable to large scale sequencing dataset, we could expand upon this further to show that this association is stronger in samples from patients with worst prognosis. This leads to the hypothesis that quantifying immune:neoplastic cell interactions could be prognostic, or that targeting them could be therapeutically beneficial, exemplifying the value in applying GBMdeconvoluteR to gain biological and clinical insights.

In summary, GBMdeconvoluteR can be used to assess associations between cell type quantities and phenotypic, molecular or clinical characteristics with applications for target identification, gaining mechanistic insight or stratifying samples for retrospective therapeutic evaluation or prospective precision medicine approaches.

## Acknowledgements

This work was supported by grants from UK Research and Innovation [MR/T020504/1 to LFS], the Integrated Biological Imaging Network [IBIN4LS to LFS], Yorkshire’s Brain Tumour Charity and OSCARs Paediatric Brain Tumour Charity [Joint Infrastructure funding to LFS], the British Neuropathology Society [Small Grant Award to SA], and Health Data Research UK, an initiative funded by UK Research and Innovation, Department of Health and Social Care (England) and the devolved administrations, and leading medical research charities. Work in the Ihrie lab is supported by the US National Institutes of Health [R01NS118580 and a supplement to U54CA217450 to RAI], the Ben & Catherine Ivy Foundation [RAI], and a gift from the Michael David Greene Brain Cancer Fund at the Vanderbilt–Ingram Cancer Center [RAI]. Tissue used in this study was accessed from the Sidney Driscol Neuroscience Foundation BTNW tissue bank. Tissue processing was possible through the Leeds Neuropathology Research Tissue Bank funded by Yorkshire’s Brain Tumour Charity and OSCARs Paediatric Brain Tumour Charity.

