## Supplemental Figures for "GBMdeconvoluteR accurately infers proportions of neoplastic and immune cell populations from bulk glioblastoma transcriptomics data"

### Slide 1
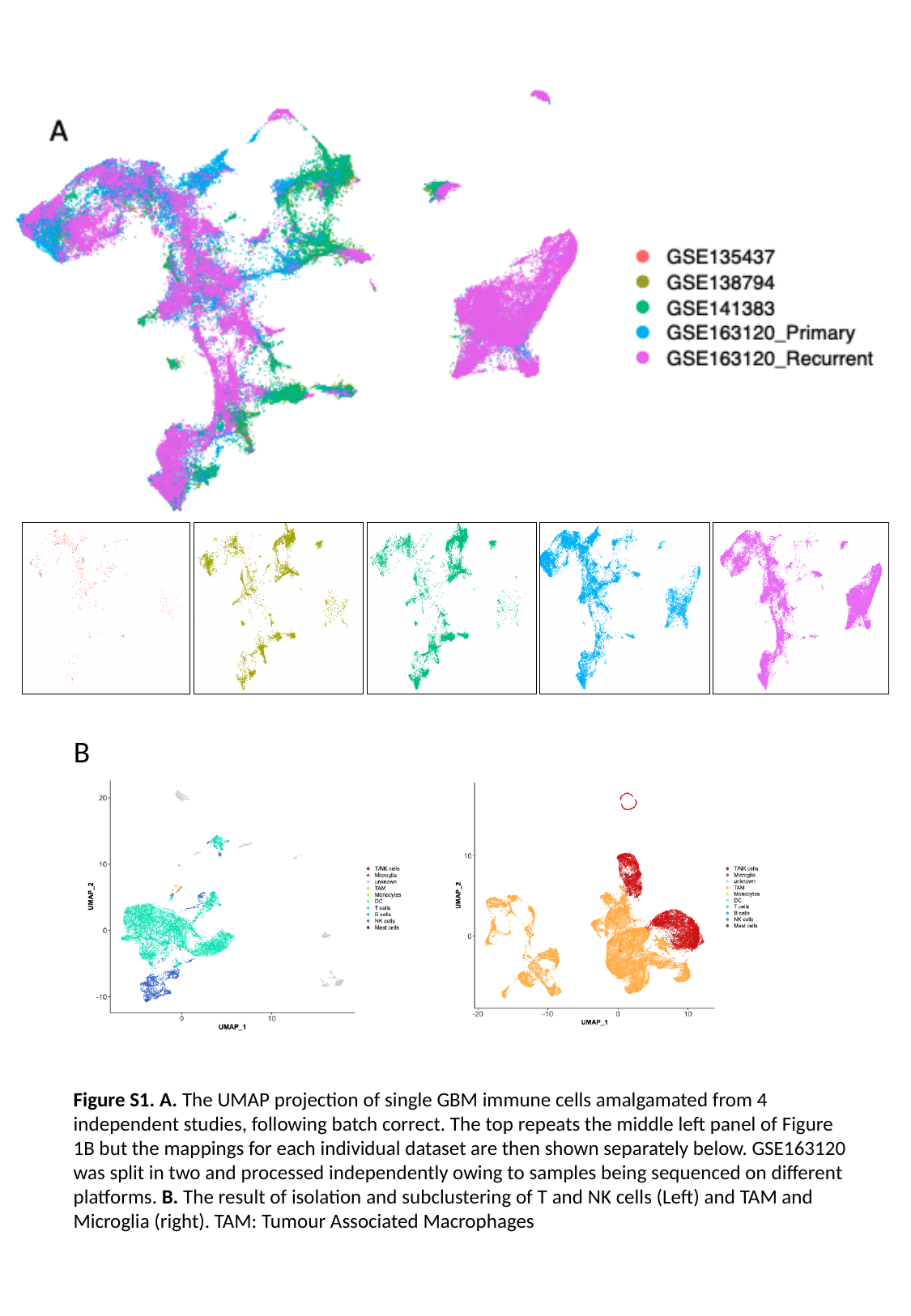

B
Figure S1. A. The UMAP projection of single GBM immune cells amalgamated from 4 independent studies, following batch correct. The top repeats the middle left panel of Figure 1B but the mappings for each individual dataset are then shown separately below. GSE163120 was split in two and processed independently owing to samples being sequenced on different platforms. B. The result of isolation and subclustering of T and NK cells (Left) and TAM and Microglia (right). TAM: Tumour Associated Macrophages

### Slide 2
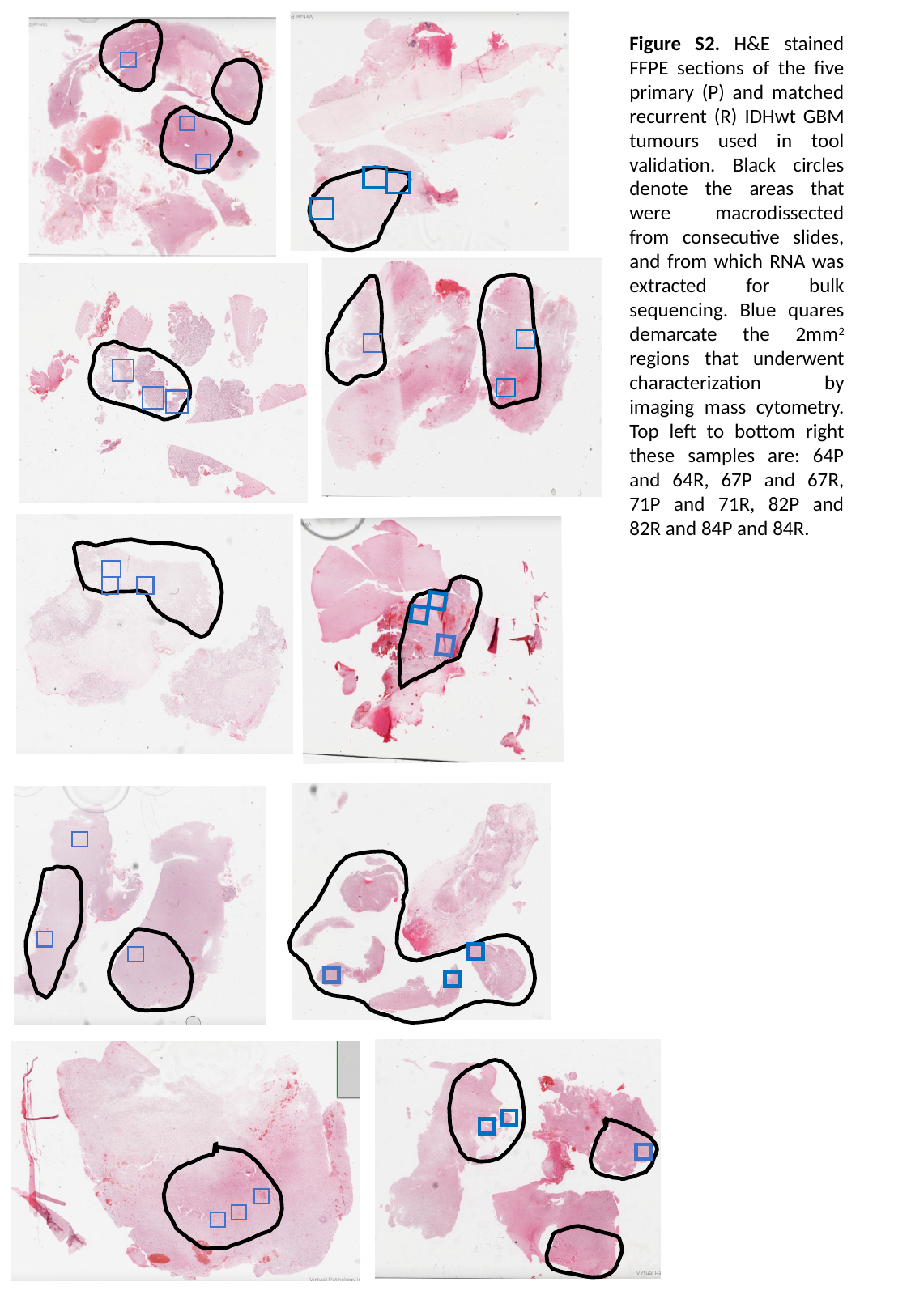

Figure S2. H&E stained FFPE sections of the five primary (P) and matched recurrent (R) IDHwt GBM tumours used in tool validation. Black circles denote the areas that were macrodissected from consecutive slides, and from which RNA was extracted for bulk sequencing. Blue quares demarcate the 2mm^2^ regions that underwent characterization by imaging mass cytometry. Top left to bottom right these samples are: 64P and 64R, 67P and 67R, 71P and 71R, 82P and 82R and 84P and 84R.
